# Insecticide susceptibility of *Aedes (Stegomyia) aegypti* (Linnaeus, 1762) and *Aedes (Stegomyia) albopictus* (Skuse, 1894) in Kinshasa, Democratic Republic of the Congo

**DOI:** 10.1101/2021.11.08.467678

**Authors:** Fabien Vulu, Gillon Ilombe, Lucrecia Vizcaino, Joachim Mariën, Yasue Morimoto, David Weetman, Audrey Lenhart, Seth R. Irish, Thierry L. Bobanga

**Affiliations:** Services de Parasitologie et d’Entomologie, Département de Médecine Tropicale, Faculté de Médecine, Université de Kinshasa, Democratic Republic of the Congo; Graduate School of Biomedical Sciences, Nagasaki University, Nagasaki, Japan; Department of Entomology, Institut National de Recherche Biomédicale, Kinshasa, Democratic Republic of the Congo; Antwerp University, Global Health Institute, Antwerp, Belgium; Entomology Branch, Division of Parasitic Diseases and Malaria, Center for Global Health, Centers for Disease Control and Prevention, Atlanta, Georgia, U.S.A; Evolutionary Ecology group, University of Antwerp, Antwerp, Belgium; Department of Vector Biology, Liverpool School of Tropical Medicine, Pembroke Place, Liverpool, United Kingdom; President’s Malaria Initiative, Centers for Disease Control and Prevention, Atlanta, Georgia, USA

**Keywords:** *Aedes* mosquitoes, pyrethroids, resistance, *kdr* mutations

## Abstract

*Aedes aegypti* and *Aedes albopictus* are arbovirus vectors of public health concern. Although the Democratic Republic of the Congo (DRC) faces a long-standing risk of *Aedes*-borne viruses, data on insecticide resistance of *Aedes* populations are absent. To address this gap, we investigated insecticide susceptibility of *Ae. aegypti* and *Ae. albopictus* in areas with a high risk of arbovirus transmission. We also investigated the frequency of knock-down resistance (*kdr*) mutations in *Ae. aegypti*. Immature stages of *Ae. aegypti* and *Ae. albopictus* were collected from two sites in Kinshasa (Lingwala and Cité Verte) between April and July 2017 and reared to the adult stage. Wild-caught adult *Ae. aegypti* were collected in 2016 in another site (Ngaliema). Female *Ae. aegypti* (from Lingwala) and *Ae. albopictus* (from Cité Verte) were used in WHO tube insecticide susceptibility tests. The F1534C, V1016I and V410L *kdr* mutations were genotyped in *Ae. aegypti* from Lingwala and Ngaliema. We observed *Ae. aegypti* to be susceptible to bendiocarb, propoxur and malathion, suspected resistant to permethrin, and resistant to deltamethrin and DDT. *Aedes albopictus* was susceptible to bendiocarb, propoxur, malathion and permethrin, suspected resistant to deltamethrin and resistant to DDT. While F1534C and V1016I were not detected, a few *Ae. aegypti* from Lingwala were heterozygous for the mutation V410L. This study reports for the first time the insecticide resistance status of *Aedes spp*. and the detection of the *kdr* mutation V410L in *Ae. aegypti* in DRC. Given the resistance profile, carbamates and potentially malathion are recommended insecticide options against *Ae. aegypti* in Kinshasa. It will be important to develop *Aedes* control strategies based on the resistance patterns of *Aedes* in Kinshasa.

## Introduction

*Aedes aegypti* and *Aedes albopictus* are important arbovirus vectors with an increasing distribution range [1]. Whereas *Ae. aegypti* originated in sub-Saharan Africa, *Ae. albopictus* originated in southeast Asia and was first observed in Africa in 1989 (South-Africa) [2, 3, 4]. Currently, *Ae. aegypti* is distributed primarily in tropical and subtropical regions of the world while *Ae. albopictus* is found on all continents except Antarctica [1, 5]. Both *Ae. aegypti* and *Ae. albopictus* can transmit viruses that are pathogenic to humans including dengue, chikungunya, Zika and yellow fever viruses [5].

The public health threat posed by *Aedes*-borne viruses is increasing due to the rapid spread of their mosquito vectors in new regions [1, 5, 6]. Dengue fever (DF) was first reported in the Americas in the seventeenth century following the introduction of *Ae. aegypti* [7]. Currently, dengue virus (DENV, *Flaviviridae, Flavivirus*) is the most important arbovirus worldwide: endemic in more than 125 countries, dengue affects an estimated 390 million people annually [8]. Chikungunya (CHIKV, *Togaviridae, Alphavirus*) and Zika (ZIKV, *Flaviviridae, Flavivirus*) viruses both emerged in East Africa in the middle of the twentieth century and then spread worldwide [9, 10]. Although both diseases are seldom deadly, they can cause a high disease burden in the affected communities [9-11]. Yellow fever (YF) originated in Africa and emerged in the Americas in the seventeenth century. Despite the existence of an effective vaccine, YF remains a public health concern in Africa and South America. It was estimated that 97,400 YF cases (28,000-251,700) and 4,800 (1,000-13.800) associated deaths occurred worldwide in 2017 [12]. The 2015-2016 YF outbreak in Angola and the Democratic Republic of the Congo (DRC) highlighted once again how threatening this disease can be [13].

In the absence of effective vaccines or specific drugs against most *Aedes*-borne viruses, vector control remains indispensable to control and prevent disease outbreaks [14]. Vector control is often based on source reduction (e.g. removing water containers that serve as larval habitats) and the use of insecticides against adult and immature mosquitoes. The insecticides generally used belong to four main families: organochlorines, organophosphates, carbamates and pyrethroids [15]. Pyrethroids are most commonly used against adult *Aedes* because of their broad arthropod toxicity but low toxicity to mammals [15, 16]. Since *Aedes* are increasingly reported as resistant to pyrethroids and other insecticides in many parts of the world, it is essential to monitor the susceptibility of *Aedes* to these insecticides in areas threatened by *Aedes*-borne viruses [14, 17]. Moreover, the determination of the underlying resistance mechanisms is of interest since it can guide the choice of new insecticides by providing a deeper understanding of cross resistance and selection pressures on populations. Insensitivity of insecticide target sites due to mutations and increased insecticide detoxification are the main mechanisms associated with resistance in *Aedes* [17]. Mutations in the voltage gated sodium channel gene causing knock-down resistance (*kdr*) are important target site mutations associated with pyrethroid resistance and are common in *Ae. aegypti* [17]. More than 10 mutations have been recorded globally, of which the F1534C, V1016I, and V410L mutations have been reported so far in *Ae. aegypti* in Africa [17-21].

Much of DRC is at high risk of *Aedes*-borne virus transmission [5, 22-25], with YF and chikungunya outbreaks occurring regularly [24 - 28]. Although DF outbreaks have not been reported, seroprevalence studies indicate that DENV strains are circulating in the human population [29 - 31]. Both *Ae. aegypti* and *Ae. albopictus* are present in DRC and large areas of the country are suitable for their establishment [5, 32]. In addition, a previous study reported that CHIKV was detected in *Aedes* populations around Kinshasa [33]. Also, *Ae. aegypti* and *Ae. albopictus* were vectors of the 2019 chikungunya outbreak in DRC’s capital Kinshasa and the major port city of Matadi, respectively [24]. Despite the long-standing risk of *Aedes*-borne diseases, insecticide resistance status of *Aedes* populations have not previously been reported from DRC [17]. To address this gap, we investigated the susceptibility of *Ae. aegypti* and *Ae. albopictus* in high risk areas of arbovirus transmission to the following insecticides: dichlorodiphenyltrichloroethane (DDT), deltamethrin, permethrin, bendiocarb, propoxur and malathion. We also investigated the frequency of three *kdr* mutations (F1543C, V1016I, V410L) in *Ae. aegypti*.

## Materials and methods

### Study sites

Immature stages of *Ae. aegypti* and *Ae. albopictus* were collected from two sites in Kinshasa (DRC) between April and July 2017, Lingwala (S 04^°^ 19’33’’, E 015^°^18’20”) and Cité Verte (S 04^°^25’52”, E 015°15’35”) (Fig. 1). An additional sample of dead adult *Ae. aegypti* collected in February 2016 in Ngaliema (S 04° 21’7’’/E 015° 14’33’’) during a previous study was also used for *kdr* genotyping only (Fig. 1) [32]. All these sites have experienced arbovirus outbreaks in the past. Autochthonous cases of YF were reported during the 2016 outbreak in Lingwala and in Selembao, in which the Cité Verte site is located. Probable chikungunya cases were observed in Cité Verte and Ngaliema during the poorly documented chikungunya outbreak in 2012 [27, 30, 31, 34]. Furthermore, antibodies against CHIKV and DENV were reported in approximately 30% of febrile patients from Selembao and Ngaliema, tested in 2005-2006 [29].

**Fig 1.**
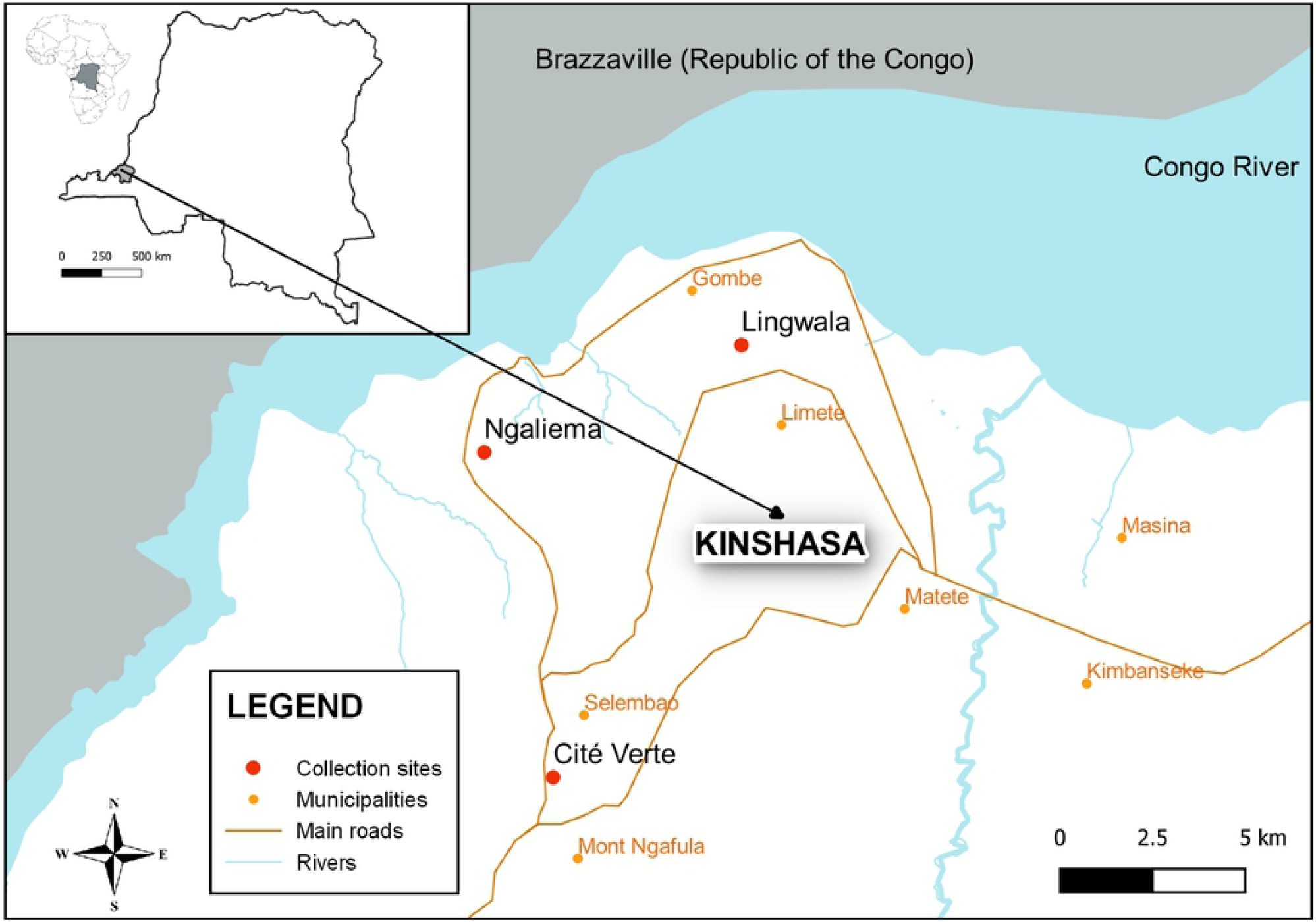
*Aedes* sampling sites in Kinshasa, DRC.

### Mosquito collections, rearing and identification

Tires filled with rain water were used as oviposition traps in sites in Lingwala (n=6) and Cité Verte (n=10). The tires were checked every five days throughout the study duration for the presence of mosquito larvae, which were collected using a pipette or ladle and brought to the insectary at the Tropical Medicine Department of Kinshasa University. Larvae were fed with fish food and reared to the adult stage under ambient conditions, with temperature and humidity within the target ranges of 28±2 °C and 80±5%, respectively, throughout the study duration. Adult mosquitoes obtained from immature stages were identified to species using morphological keys according to Huang [35] and kept alive. In the earlier sampling in Ngaliema, adult mosquitoes were collected using electric Prokopack aspirators (John W. Hock, Gainesville, USA) from 3:30 to 6:30 pm [32]. The dead mosquitoes were later identified to species as above.

### Adult bioassays

To assess the insecticide susceptibility of *Ae. aegypti* and *Ae. albopictus*, we used World Health Organisation (WHO) insecticide susceptibility tube tests performed on (non-blood-fed) F0 adult female *Ae. aegypti* (from Lingwala) and *Ae. albopictus* (from Cité Verte) [36]. For each test, 3-4 replicates of 20-25 females fed *ad libitum* on a 10% sugar solution were exposed for one hour to insecticide-impregnated papers treated with deltamethrin 0.05% (pyrethroid), permethrin 0.75% (pyrethroid), bendiocarb 0.1% (carbamate), propoxur 0.1% (carbamate), malathion 5% (organophosphate), or DDT 4% (organochlorine). Although these insecticide concentrations were initially recommended for *Anopheles*, they are commonly used to screen *Aedes* for insecticide resistance in Africa, though it should be noted that the doses for permethrin and malathion are three and approximately six-times higher, respectively, than the recommended *Aedes* doses [17]. Insecticide-impregnated papers were obtained from the Vector Control Research Unit at the Universiti Sains Malaysia, Penang, Malaysia. Controls were also run by exposing 2 replicates of 20-25 adult female mosquitoes to untreated papers. After 60 minutes of insecticide exposure, mosquitoes were transferred into holding tubes and supplied with a 10% sugar solution. Percent mortality was recorded 24 hours after exposure, and the 95% binomial confidence intervals were calculated using SPSS 21.0 (IBM Corp. Armonk, NY, USA).

Mortalities were corrected when necessary using Abbott’s formula (if control mortality was between 5 and 20%) [36]. We used the following WHO criteria to score population-level resistance/susceptibility: Mortality rates <90% were considered resistant, > 97% were considered susceptible and mortality rates between 90 and 97% were suspected resistant [36]. *Ae. aegypti* specimens that survived after pyrethroid exposures were killed by freezing, stored at 0°C and later sent (with *Ae. aegypti* from Ngaliema) to the Centers for Disease Control and Prevention (CDC, USA) for *kdr* genotyping.

### DNA extraction and *kdr* mutation detection

Real-time PCR was used to identify the F1534C, V1016I, and V410L *kdr* mutations. To estimate the allele frequencies, 28 *Ae. aegypti* female survivors to pyrethroid exposures from Lingwala and 47 wild caught female *Ae. aegypti* from Ngaliema were analyzed. DNA was extracted from individual mosquitoes using the Quanta Biosciences ExtractaTM Kit. Each mosquito was placed in a sterile 0.2 mL tubes with 25 μL extraction buffer, followed by an incubation at 95°C for 30 min in a C1000 Bio-Rad CFX 96 TouchTM Real-Time System thermocycler. At the end of the incubation, 25 μL of stabilization buffer was added. DNA was quantified using a NanoDropTM 2000/2000c spectrophotometer (ThermoFisher Scientific). PCR reactions were performed in a Bio-Rad C1000 CFX96 Real-Time System thermocycler. Genotypes were determined by analyzing the melting curves of the PCR products.

The F1534C mutation was detected following the methodology described by Yanola et al. [37] using a final reaction volume of 20 μL comprised of 7.15 μL of ddH2O, 9 μL of iQTM SYBR1Green Supermix (Bio-Rad), 0.6 μL of each of the F1534-f forward primer, [5’-GCG GGC TCT ACT TTG TGT TCT TCA TCA TAT T-3’] and CP-r common reverse primer, [5’-TCT GCT CGT TGA AGT TGT CGA T-3’]; 0.65 μL of the C1534-f forward primer, [5’-GCG GGC AGG GCG GCG GGG GCG GGG CCT CTA CTT TGT GTT CTT CAT CAT GTG-3’] primer, and 2 μL of DNA template. The cycling conditions were as follows: an initial denaturation at 95°C for 3 min followed by 37 cycles of: 95°C for 10 s, 57°C for 10 s, and 72°C for 30 s; and a final extension at 95°C for 10 s. The melting curves were determined by a denaturation gradient from 65°C to 95°C with an increase of 0.2°C every 10 s. The V1016I mutation was amplified following the methodology described by Saavedra-Rodriguez *et al*. [38], using a final reaction volume of 20 μL, containing 8.866 μL of ddH2O, 8 μL of iQTM SYBR1 Green Supermix (Bio-Rad), 0.4 μL of each of the Iso1016f forward primer, [5’-GCG GGC ACA AAT TGT TTC CCA CCC GCA CTG A-3’]; and Iso1011r reverse primer ; [5’-GGA TGA ACC SAA ATT GGA CAA AAG C-3’]; 0.34 μL of Val1016f forward primer [5’-GCG GGC AGG GCG GCG GGG GCG GGG CCA CAA ATT GTT TCC CAC CCG CAC CGG-3’] and 1 μL of DNA template. The cycling conditions were as follows: an initial denaturation at 95°C for 3 min followed by 35 cycles of: 95°C for 10 s, 60°C for 10 s, and 72°C for 30 s; and a final extension at 95°C for 10 s.

The melting curves were determined by a denaturation gradient from 65°C to 95°C with an increase of 0.2 °C every 10 s.

The V410L mutation was detected following the methodology described by Saavedra *et al*. [39] using a final reaction volume of 20 μL comprised of 8.7 μL of ddH2O, 9.9 μL of iQTM SYBR1Green Supermix (Bio-Rad), 0.1 μL of each L410fw, [5’-GCG GGC ATC TTC TTG GGT TCG TTC TAC CAT T-3’] and V410fw, [5’-GCG GGC AGG GCG GCG GGG GCG GGG CCA TCT TCT TGG GTT CGT TCT ACC GTG-3’] primers; 0.2 μL of a common reverse primer 410rev [5’-TTC TTC CTC GGC GGC CTC TT-3’] and 2 μL of DNA template. The cycling conditions were as follows: an initial denaturation at 95°C for 3 min followed by 40 cycles of: 95°C for 10 s, 60°C for 10 s, and 72°C for 30 s; and a final extension at 95°C for 10 s. The melting curves were determined by a denaturation gradient from 65°C to 95°C with an increase of 0.2°C every 10 s.

## Results

### Relative abundance of *Aedes* species by collection site

A total of 4,802 *Aedes* were obtained from immature stages collected throughout the study belonging to two species: *Ae. aegypti* (2,558 specimens) and *Ae. albopictus* (2,244 specimens). A total of 2,544 (1,576 females) *Ae. aegypti* and 171 (93 females) *Ae. albopictus* were obtained from Lingwala and 14 (4 females) *Ae. aegypti* and 2,073 (1,320 females) *Ae. albopictus* were obtained from Cité Verte. In total 1,851 female *Aedes* were used in bioassays (including controls and 2 tests discarded because of high mortality in the control), out of which 1,097 were female *Ae. aegypti* from Lingwala and 754 were female *Ae. albopictus* from Cité Verte.

### Adult bioassays

*Aedes aegypti* were fully susceptible to bendiocarb and malathion (100% mortality) and also susceptible to propoxur (98% mortality). Suspected resistance was detected to permethrin (97% mortality), and resistance was detected to deltamethrin and to DDT with mortality rates of 73% and 25%, respectively (Table 1). *Aedes albopictus* was fully susceptible to permethrin, bendiocarb, propoxur and malathion (100% mortality). Suspected resistance to deltamethrin (92% mortality) and resistance to DDT (36% mortality) were detected (Table 1).

**Table 1:** Mortality rates of adult female *Ae. aegypti* (Lingwala) and *Ae. albopictus* (Cité Verte) 24 hours after exposure to insecticides in WHO bioassays.

| Insecticide class | Insecticide type | *Aedes* species | Number tested<br>(mosquito dead) | % Mortality<br>(95% CI) | Status <sup>a</sup> |
| --- | --- | --- | --- | --- | --- |
| Organochlorine | DDT 4% | *Ae. aegypti* | 63 (16) | 25 (16 - 37) | **R** |
|  |  | *Ae. albopictus* | 84 (30) | 36 (26 - 46) | **R** |
| Pyrethroid | Deltamethrin 0.05% | *Ae. aegypti* | 94 (69) | 73 (64 - 82) | **R** |
|  |  | *Ae. albopictus* | 80 (74) | 92 (86-98) | **SR** |
|  | Permethrin 0.75% | *Ae. aegypti* | 97 (94) | 97 (93 - 100) | **SR** |
|  |  | *Ae. albopictus* | 80 (80) | 100 (95-100) | **S** |
| Carbamate | Bendiocarb 0.1% | *Ae. aegypti* | 93 (93) | 100 (96-100) | **S** |
|  |  | *Ae. albopictus* | 80 (80) | 100 (95-100) | **S** |
|  | Propoxur 0.1% | *Ae. aegypti* | 98 (96) | 98 (95 - 100) | **S** |
|  |  | *Ae. albopictus* | 80 (80) | 100 (95-100) | **S** |
| Organophosphate | Malathion 5% | *Ae. aegypti* | 94 (94) | 100 (96-100) | **S** |
|  |  | *Ae. albopictus* | 100 (100) | 100 (96-100) | **S** |

<sup>a</sup> R: Resistant; S: Susceptible; SR: Suspected Resistant.

### *kdr* mutations

A total of 75 female *Ae. aegypti* (28 from Lingwala and 47 from Ngaliema) were genotyped for the *kdr* mutations F1534C and V1016I. All mosquitoes tested were wild type at these loci. Concerning mutation V410L, 54 female *Ae. aegypti* (24 from Lingwala and 30 from Ngaliema) were genotyped. Seven mosquitoes were heterozygous for the leucine mutation (all from Lingwala) with a resulting frequency of L410 at 14.5% (CI 95%: 8.7-20.3%) (Table 2).

**Table 2:** Frequency of V410L *kdr* mutation in *Ae. aegypti*.

| Site | n | 410 |  |  | L410 Allele frequency |  |
| --- | --- | --- | --- | --- | --- | --- |
|  |  | VV | VL | LL | Frequency (%) | 95% CI |
| Lingwala | 24 | 17 | 7 | 0 | 14.5 | 8.7–20.3 |
| Ngaliema | 30 | 30 | 0 | 0 | 0 | – |

## Discussion

This study determined the susceptibility of *Ae. aegypti* and *Ae. albopictus* from Kinshasa to insecticides. Both *Ae. aegypti* and *Ae. albopictus* showed a high frequency of resistance to DDT and moderate resistance to pyrethroids, although were susceptible to carbamates and organophosphates. These results are consistent with other studies performed in Africa. Results from bioassays in neighbouring Republic of the Congo (RC) and Central African Republic (CAR) showed populations of *Ae. aegypti* and *Ae. albopictus* highly resistant to DDT [40, 41]. Highly-resistant populations to DDT were also recorded elsewhere in Africa including Cameroon and Burkina Faso [42, 43], as well as in Asia and in the Americas [17, 44 - 46]. Some authors have explained the high frequency of DDT resistance in *Aedes* populations as being due to the intense use of this insecticide in the past [40, 41]. Indeed, the use of DDT in aerial spraying to control malaria vectors in Kinshasa several decades ago and the intense use in farming could have led to the emergence of resistance in mosquitoes including *Aedes spp*. [47]. Considering the pyrethroids, *Ae. aegypti* was resistant to deltamethrin and suspected resistant to permethrin, while *Ae. albopictus* was suspected resistant to deltamethrin and susceptible to permethrin. Fully susceptible populations of both species to deltamethrin have been recorded in RC, Cameroon, CAR and Nigeria; however, suspected or moderately resistant populations were also recorded by the same studies [40 - 42, 48]. Although, *Ae. aegypti* and *Ae. albopictus* populations have generally shown low levels of resistance to deltamethrin in other parts of Africa, studies in Burkina Faso detected the presence of highly resistant *Ae. aegypti* populations [43, 49, 50]. Also, in Asia and in the Americas, studies have reported the presence of *Aedes* populations highly resistant to deltamethrin [17]. Regarding permethrin, studies in RC and Cameroon detected *Ae. aegypti* and *Ae. albopictus* resistant populations using the insecticide concentration that WHO typically recommends for *Aedes* [41, 42]. However, our study used a permethrin concentration three-fold higher [51]. As such, the permethrin resistance level could have been underestimated. Nevertheless, resistant *Ae. aegypti* and *Ae. albopictus* populations were also recorded in studies in Burkina Faso, Tanzania and Ghana using permethrin concentration similar to our study [52 - 54]. The reduced susceptibility to pyrethroids in both *Ae. aegypti* and *Ae. albopictus* observed in this study is of concern as pyrethroids are broadly recommended in control activities against adult *Aedes* [15, 16]. Moreover, the intense and continuous use of pyrethroids in mosquito nets, home insecticide sprays, and farming in Kinshasa might exacerbate the mosquito resistance [55].

Both *Ae. aegypti* and *Ae. albopictus* were susceptible to bendiocarb, propoxur and malathion. These results are similar to other studies performed in neighbouring countries including RC, CAR and Tanzania in which some *Aedes* populations were susceptible to carbamates and organophosphates [40, 41, 56]. In other parts of Africa (e.g. Burkina Faso), carbamate-resistant *Aedes* populations are more common [49, 50]. On the other hand, bioassays conducted with malathion across Africa have shown that both *Ae. aegypti* and *Ae. albopictus* populations were susceptible to this insecticide [34, 40, 48 - 50, 53]. This result is in contrast to some *Aedes* populations observed in the Americas and Asia where high malathion resistance has been reported [17, 45, 57]. However, we highlight that our study (and most other studies performed in Africa) used a malathion concentration that was six times higher than what is typically recommended for *Aedes* by the WHO [51]. Our results suggest that carbamates and potentially malathion could be recommended insecticide options against *Ae. aegypti* in Kinshasa.

To determine the underlying resistance mechanisms in *Ae. aegypti*, we also tested for *kdr* mutations. While the *kdr* mutations F1534C and V1016I were not detected in our study, V410L was detected at a low frequency in the *Ae. aegypti* population from Lingwala. The F1534C mutation was previously reported in *Ae. aegypti* in West Africa [21, 49, 53, 58] but not in three studies conducted in central Africa [19, 40, 41]. The V1016I mutation in *Ae. aegypti* was also reported in Africa in Ghana, Burkina Faso, Cote d’Ivoire, Angola and Cape Verde [18, 19, 21, 49]. The V410L mutation was first detected in *Ae. aegypti* in Brazil [59]. Results from that study [59] revealed that this mutation alone could be responsible for a decrease in *Ae. aegypti* susceptibility to insecticides. Another study in Mexico reported this mutation in samples of *Ae. aegypti* collected in 2002 [39], well before the samples from the Brazilian study. The Mexican study revealed an increase in the frequency of this mutation reaching very high levels by 2016 [39]. In Africa, the V410L mutation was first reported in *Ae. aegypti* in Angola and Cape Verde [19]. This low mutation frequency was surprising given the usual tight linkage observed between the V1016I and V410L *kdr* loci, although occurrence of the 410L mutation alone has occasionally been observed [19, 39]. While mosquito surviving pyrethroid exposures were used for *kdr* genotyping in *Ae. aegypti* from Lingwala, mosquitoes of unknown resistance phenotype were used from Ngaliema. The V410L mutation may be contributing to *Aedes* insecticide resistance in Kinshasa, but given its low frequency, there are likely other resistance mechanisms that are also important. Future research can hopefully explore what those mechanisms are, as well as the frequencies of *kdr* mutations and other mechanisms to mutations that might cause resistance in *Ae. albopictus*.

This study had several limitations. First, the insecticide susceptibility tests were performed on mosquitoes from limited sampling areas. Moreover, higher doses of permethrin and malathion than recommended for *Aedes* were used in the bioassays, so resistance may be underestimated. Also, a limited number of mosquitoes were used to detect *kdr* mutations, so sampling a greater proportion of mosquitoes would give a more accurate estimate of allele frequencies. Despite these limitations, this study is valuable because for the first time, it reports data on the resistance patterns of *Aedes* in Kinshasa, including the detection of the *kdr* mutation V410L in *Ae. aegypti* in the DRC. This information will be useful to guide future insecticide resistance surveillance programmes and to develop control strategies for areas at high risk of *Aedes*-borne arboviruses in the DRC.

### Disclaimer

The views expressed in this manuscript are those of the authors and do not necessarily reflect the official policy or position of the U.S. Centers for Disease Control and Prevention.

## Acknowledgments

The authors would like to thank Vestergaard for the help to obtain insecticide impregnated paper.

## Supporting file captions

**S1 Appendix. Frequency of *kdr* mutations in *Ae. aegypti***.

